# NDRG2 Regulates Adherens Junction Integrity to Restrict Colitis and Tumorigenesis

**DOI:** 10.1101/473397

**Authors:** Mengying Wei, Yongzheng Ma, Liangliang Shen, Yuqiao Xu, Lijun Liu, Xin Bu, Zhihao Guo, Hongyan Qin, Zengshan Li, Zhe Wang, Kaichun Wu, Libo Yao, Jipeng Li, Jian Zhang

## Abstract

Paracellular barriers play an important role in the pathogenesis of IBDs and maintain gut homeostasis. N-myc downstream-regulated gene 2 (NDRG2) has been reported to be a tumor suppressor gene and inhibits colorectal cancer metastasis. However, whether NDRG2 affects colitis initiation and colitis-associated colorectal cancer is unclear. Here, We found that intestine-specific *Ndrg2* deficiency caused mild spontaneous colitis with ageing, aggravated DSS and TNBS induced colitis, increased AOM-DSS induced colitis-associated tumor. *Ndrg2* loss led to adherens junction (AJ) structure destruction *via* E-cadherin expression attenuation, resulting in diminished epithelial barrier function and increased intestinal epithelial permeability. Mechanistically, NDRG2 enhancing the interaction of E3 ligase FBXO11 with Snail, the repressor of E-cadherin, to promote Snail degradation by ubiquitination, and maintained E-cadherin expression. In human ulcerative colitis patients, reduced NDRG2 expression is positively correlated with severe inflammation. These findings demonstrate that NDRG2 is an essential colonic epithelial barrier regulator and plays important role in gut homeostasis maintenance and colitis-associated tumor development.

**SUMMARY:** Adherens junctoin (AJ) as the key part of intestinal epithelial barrier plays important role in the pathogenesis of IBDs. Intestinal specific *Ndrg2* loss attenuates E-cadherin expression and disrupts the integrity of AJ structure which is feasible for colitis and tumor development.

## INTRODUCTION

Inflammatory bowel disease (IBD) is a multifactorial chronic inflammatory disease mainly comprising Crohn’s disease (CD) and ulcerative colitis (UC) that causes intestinal epithelial cell injury and relapsing chronic pathogenic rectal and colonic inflammation(Arrieta et al., 2006; de Souza et al., 2017). Intestinal epithelial homeostasis plays an important role in the intestinal tract. This bacteria-host homeostasis is maintained by a epithelial barrier, which includes tight junctions (TJs), adherens junctions (AJs) and desmosomes(Stein et al., 1998; Taddei et al., 2008). AJs are cell-cell adhesion complexes that participate in embryogenesis and tissue homeostasis(Smalley-Freed et al., 2010). The major AJ component E-cadherin or p120-catenin loss results in embryonic death(Arrieta et al., 2006). Previous studies demonstrated that AJ dysfunction is associated with IBD and E-cadherin expression attenuation could obviously aggravate colitis, possibly due to increase of colonic epithelial barrier permeability(Grill et al., 2015).

We previously identified N-myc downstream regulated gene 2 (NDRG2) as a novel tumor suppressor gene that played a role in regulating the proliferation, differentiation and metastasis of multiple types of malignant tumors(Deng et al., 2003; Zhang et al., 2006). Notably, our recent data showed that NDRG2 could inhibit colorectal cancer cell proliferation and promote cell differentiation(Shen et al., 2018). Moreover, we found that decreased NDRG2 expression was a powerful and independent predictor of a poor prognosis in colorectal cancer(Chu et al., 2011; Shen et al., 2014). However, whether NDRG2 participates in intestinal epithelial homeostasis and colitis initiation remains unknown.

In this study, we generated intestine-specific conditional *Ndrg2* knockout mice and examined the roles of *Ndrg2* in AJ structure and permeability regulation in the setting of spontaneous and experimentally induced colitis. Furthermore, we characterized the detailed function and mechanism of NDRG2 in intestinal epithelial inflammation and evaluated the effects of *Ndrg2* deficiency on gut inflammation and colitis-associated tumor.

## RESULTS

### Intestinal *Ndrg2* knockout mice develops mild spontaneous colitis

To determine the role of NDRG2 in colitis initiation, we generated intestine-specific conditional *Ndrg2* knockout mice featuring intestinal epithelial cells lacking *Ndrg2* (Supplementary figure 1A). NDRG2 expression was specifically and completely abolished in the intestinal epithelial cells (IECs) of *Ndrg2^-/-^* mice (Supplementary figure 1B, C). The mice were kept in specific-pathogen-free (SPF) condition and monitored the body weight each week. Intriguingly, we found the obvious growth retardation of *Ndrg2^-/-^* mice compared with the wild type (WT) group, especially after 16 weeks age (figure 1A). However, there was no significant difference of the activity and diet between two group mice. Thus, it’s curious for us to detect the alteration of intestinal tissues. As the Haematoxylin Eosin (H&E) staining data shown in figure 1B, there was increased intestinal inflammation in *Ndrg2^-/-^* mice and more severe with ageing, which indicated the spontaneous colitis in *Ndrg2^-/-^* mice.

**Figure 1.**
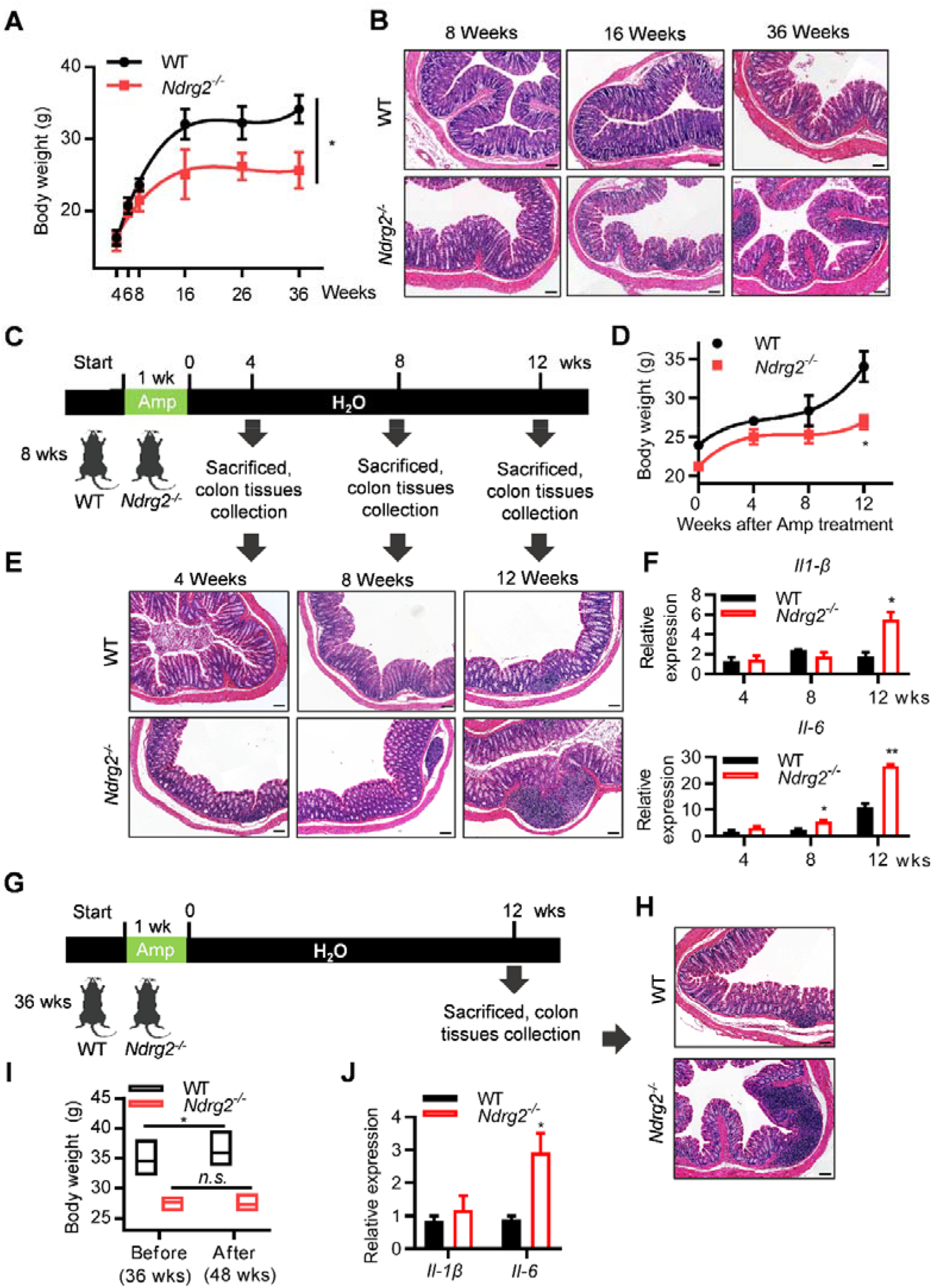
Intestine-specific conditional *Ndrg2* knockout mice (*Ndrg2^-/-^*) spontaneously developed into mild colitis with ageing. **(A)** Body weight of WT and *Ndrg2^-/-^* mice on indicated time with specific-pathogen-free (SPF) feeding condition (n=8/group; *p<0.05). **(B)** Representative H&E-staining of colon section from WT and *Ndrg2^-/-^* mice with indicated ages. **(C)** Schematic diagram for analyzing the occurrence of spontaneous colitis in 8 weeks age WT and *Ndrg2^-/-^* mice after withdrawal of ampicillin treatment for 1 week. **(D)** Body weight of WT and *Ndrg2^-/-^* mice on indicated time after ampicillin withdrawal (n=8/group; *p<0.05). **(E)** Representative H&E-staining of colon tissue. **(F)** Quantitative real-time PCR (qRT-PCR) for *Il-1β* and *Il-6* in IECs from *WT* and *Ndrg2^-/-^* mice (n=8/group; *p<0.05, **p<0.01). **(G)** Schematic diagram for analyzing the occurrence of spontaneous colitis in 36 weeks age *WT* and *Ndrg2^-/-^* mice after withdrawal of ampicillin treatment for 1 week. **(H)** Representative H&E-staining of colon section of (G). **(I)** Body weight of 36 weeks age *WT* and *Ndrg2^-/-^* mice before and after ampicillin withdrawal (48 weeks age) induced spontaneous colitis (n=8/group; *p<0.05). **(J)** qRT-PCR for *Il-1β* and *Il-6* in IECs of (G) (n=8/group; *p<0.05).

Nextly, to confirm the growth retardation of *Ndrg2^-/-^* mice was mainly due to increased intestinal inflammation, we treated the WT and *Ndrg2^-/-^* mice (8 weeks age) with ampicillin for one week to clean up the gut microbiota, and then monitored the body weight in each group (figure 1C). As anticipated, following the mice growth, there was a noticeable slower body weight increase of *Ndrg2^-/-^* mice comparing to the WT group (figure 1D). And also there was more severe intestinal inflammation and increased IL-1β and IL-6 expression in *Ndrg2^-/-^* mice (figure 1E, F), further supporting the fact of increased spontaneous colitis in *Ndrg2^-/-^* mice. Then we used the old age mice to further confirm this observation. As shown in figure 1G, both of the WT and *Ndrg2^-/-^* mice with 36 weeks age were treated with ampicillin for 1 week and analyzed the body weight and intestinal inflammation after 12 weeks respectively. Compared with the increased body weight of WT mice after 12 weeks, there was no significant alteration of *Ndrg2^-/-^* mice (figure 1I). Similar to the findings in young group mice (figure 1E, F), there was increased intestinal inflammation, IL-1β and IL-6 expression in *Ndrg2^-/-^* mice (figure 1H, J). Collectively, these data strongly suggested that intestinal *Ndrg2* knockout facilitated the development of mild spontaneous colitis.

### Intestinal *Ndrg2* loss aggravates chemical-induced colitis

To further determine the role of NDRG2 in colitis initiation and progression, we treated the mice with DSS (Dextran Sodium Sulphate) and TNBS (2, 4, 6-Trinitrobenzenesulfonic acid) respectively to mimic the colitis in vivo. We found that intestinal *Ndrg2^-/-^* mice exhibited increased body weight loss compared with WT mice with DSS treatment (figure 2A), as well as more severe colonic shortening than their counterparts (figure. 2B, C). For histological analysis, *Ndrg2^-/-^* mice exhibited severe colitis characterized by profound epithelial structural damage and robust inflammatory cell infiltration than WT group (figure 2D), as well as higher Disease Activity Index (DAI) scores (figure 2E). Alternatively, we observed the similar results with more severe inflammation of *Ndrg2^-/-^* mice in TNBS-induced colitis model (Supplementary figure 2A, B). For survival analysis, *Ndrg2^-/-^* mice exhibited increased susceptibility to lethal colitis induction (4% DSS for 7 days), leading to short survival compared with WT mice (figure 2F). Taken together, our data demonstrated that intestine-specific *Ndrg2* deficiency obviously aggravated chemical-induced colitis.

**Figure 2.**
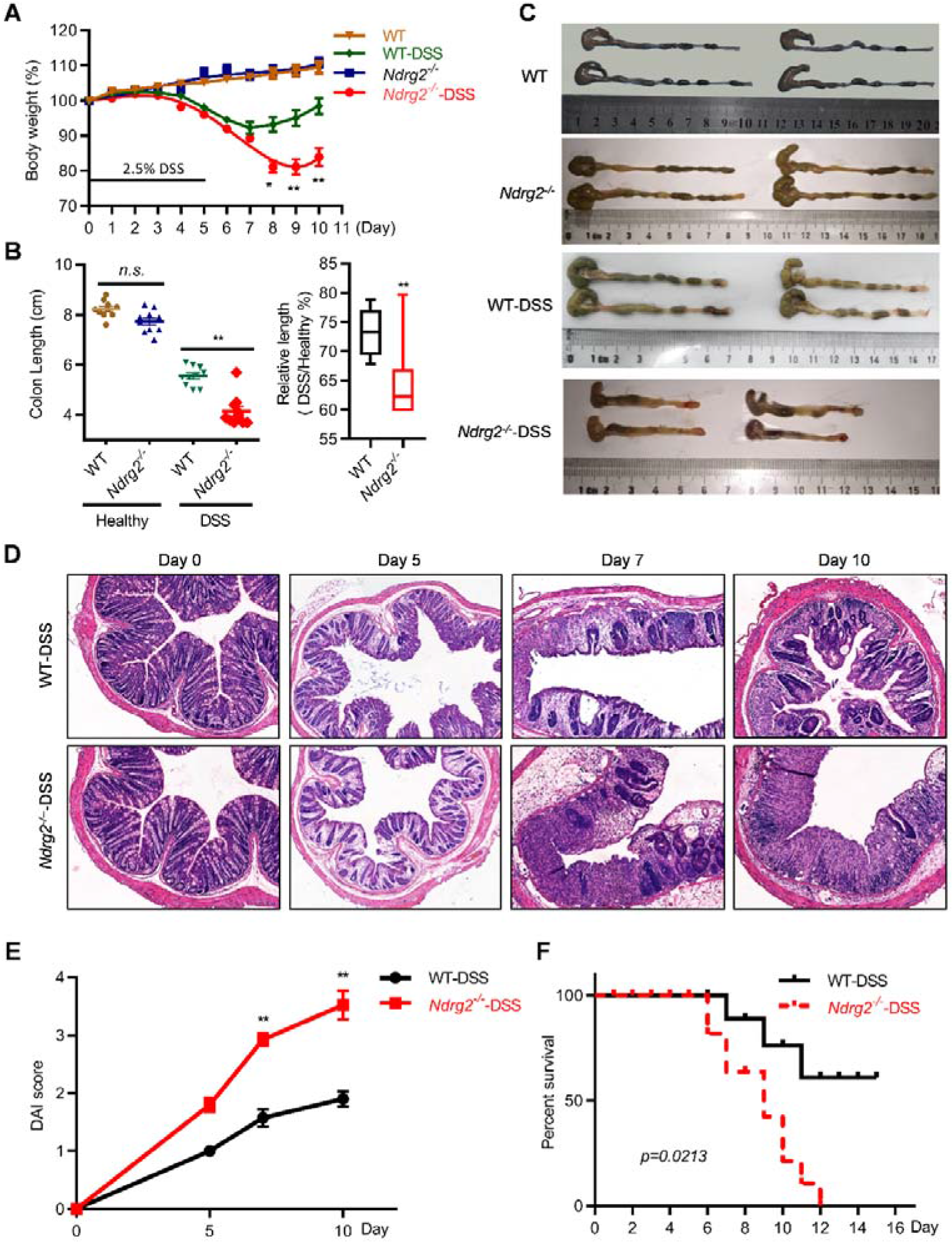
Intestine *Ndrg2^-/-^* mice exhibited increased susceptibility to dextran sodium sulfate (DSS)-induced colitis. **(A)** Body weights change of 8 weeks age WT and *Ndrg2^-/-^* mice with 2.5% DSS treatment (n=10/group; *p<0.05, **p<0.01). **(B)** Colon lengths (Left) and the statistic result (Right) on day 10 in the WT and *Ndrg2^-/-^* mice (n=10/group; **p<0.01). **(C)** Representative images of the colon appearance at the end of experiments. **(D)** Representative H&E-staining of colon tissue. **(E)** The DAI scores of the WT and *Ndrg2^-/-^* mice on days 0, 5, 7 and 10 of 2.5% DSS administration. **(F)** Survival rate analysis of 4% DSS-induced lethal colitis (n=15/group).

### Intestinal *Ndrg2* deficiency promotes inflammatory cell infiltration, chemokines and cytokines expression and LPS permeability

Inflammatory cell infiltration plays an important role in both UC and CD(Huang et al., 2019). To characterize the role of NDRG2 in colonic inflammation, IECs and lamina propria cells (LPCs) were isolated from WT and *Ndrg2^-/-^* mice treated with or without 2.5% DSS. Flow cytometry analysis showed that *Ndrg2^-/-^* mice exhibited slight increases in monocyte and macrophage infiltration compared with WT mice with DSS-free condition (figure 3A, Supplementary figure 3A). Notably, *Ndrg2^-/-^* mice exhibited significantly increased monocyte, macrophage and neutrophil infiltration with DSS treatment compared with WT mice (figure 3B, supplementary figure 3A). Furthermore, DSS-treated *Ndrg2^-/-^* mice exhibited dramatically increased the chemokines expression of CCL5 and CXCL1/2, and also pro-inflammatory cytokines expression of *IL-1β, TNF-α* and *IL-6* (figure 3C, Supplementary figure 3B). Whereas, the mRNA expression of the anti-inflammatory cytokine *IL-10* was attenuated in DSS-treated *Ndrg2^-/-^* mice (figure 3C), while the expression of *TGF-β, IL-12, IL-17 and IL-22* was comparable between the two groups (Supplementary figure 3B). These findings indicated that intestinal *Ndrg2* deficiency aggravated DSS-induced colitis by increasing chemokine and pro-inflammatory cytokine expression and promoting inflammatory cell infiltration.

**Figure 3.**
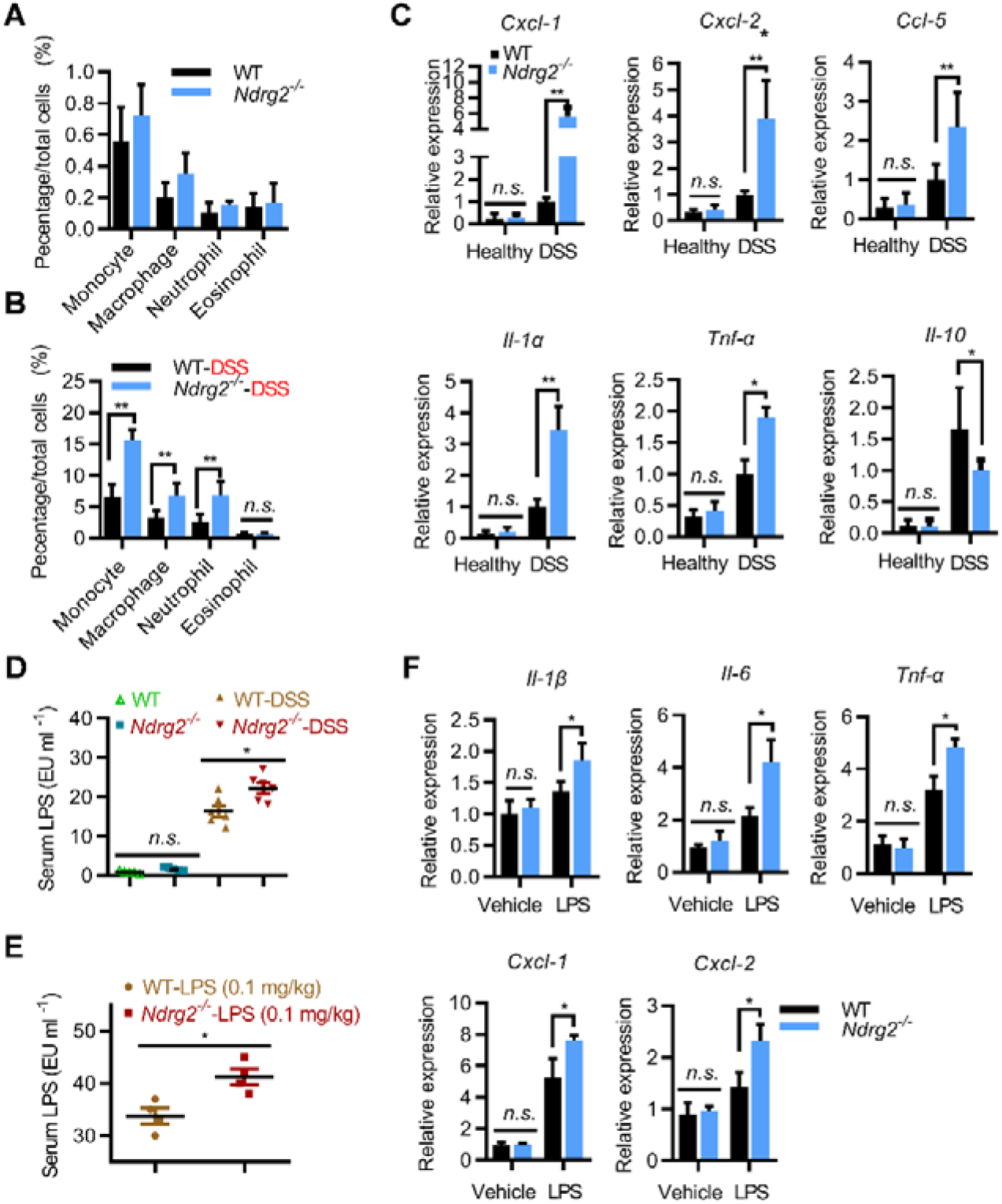
*Ndrg2* deficiency enhanced inflammation in colon characterized by increased inflammatory cell infiltration and LPS permeability. WT and *Ndrg2^-/-^* mice were treated with 2.5% DSS (n=12/group). Quantification of the infiltrated immunocytes on day 0 **(A)** and day 10 **(B)** detected by Flow cytometry analysis. **(C)** qRT-PCR analysis of chemokines *Cxcl-1, Cxcl-2, Ccl-5* and pro-inflammatory cytokines *Il-1β* and *Tnf-α*, and anti-inflammatory cytokines *Il-10*. **(D)** Detection of serum LPS concentrations in WT and *Ndrg2^-/-^* mice with and without 2.5% DSS treatment (n=6/group). **(E)** Detection of serum LPS levels in WT and *Ndrg2^-/-^* mice with 0.1 mg/kg LPS treatment with intrarectal adminstration after ampicillin treatment for 5 days, (n=4/group). **(F)** qRT-PCR analysis for *IL-1β, IL-6, TNF-α, CXCL-1 and CXCL-2* expression in IECs of WT and *Ndrg2^-/-^* mice treated with 0.1 mg/kg LPS (n=4/group). Data are represented as the mean ± SEM. *p<0.05, **p<0.01, n.s., no significance (Student’s t test).

Given that gut microbiota plays important roles in the pathogenesis of IBD and lipopolysaccharide (LPS) is the main microflora-derived toxin that induces gut epithelial inflammation. Noticeably, the serum LPS concentration is obviously increased in IBD patients(Pastorelli et al., 2015). As shown in figure 3D, DSS-treated *Ndrg2^-/-^* mice exhibited increased serum LPS levels compared with WT mice. So we subsequently examined whether *Ndrg2* deficiency facilitated LPS-induced inflammation. As expected, LPS induced significant inflammation characterized by dose-dependent increases in the levels of IL-1β, IL-6, TNF-α and CXCL1/2 expression (Supplementary figure 4 A-E). And *Ndrg2^-/-^* mice exhibited obviously higher LPS permeability and increased IL-1β, IL-6, TNF-α, CXCL-1/2 expression (figure 3E, F), suggesting the enhanced inflammatory response. These data suggested that intestine-specific *Ndrg2* deficiency increased LPS permeability and promoted LPS-induced inflammation and colitis progression.

### Intestinal *Ndrg2* loss disrupts the adherens junction integrity of normal epithelium and increased colonic permeability

Epithelial barrier dysfunction increased gut permeability and aggravated the inflammatory response in IBD. To further investigate the role of *Ndrg2* in DSS-induced colitis, we examined the gut permeability in WT and *Ndrg2^-/-^* mice treated with or without DSS *via* FITC-dextran analysis. As shown in figure 4A, even with DSS-free treatment, *Ndrg2^-/-^* mice exhibited slightly increased permeability of low molecular weight dextran. As expected, DSS-treated *Ndrg2^-/-^* mice extremely increased the permeability of both low and high molecular weight dextran, strongly suggesting epithelial barrier dysfunction in *Ndrg2^-/-^* mice. Structure disruption of adherens junction (AJ) and tight junction (TJ) resulted in epithelial barrier dysfunction, increased epithelial permeability and inflammation(Mohanan et al., 2018; Tanaka et al., 2015). Nextly, it’s much curious for us to analyze the epithelial barrier structure difference between WT and *Ndrg2^-/-^* mice. With electron microscopy analysis, we noticed that *Ndrg2^-/-^* mice exhibited significant AJ structural disruption but virtually no TJ structural alteration compared with WT mice (figure 4B). The AJ distance were remarkably increased in the intestinal epithelium of *Ndrg2^-/-^* mice (figure 4C), indicating intestinal epithelial AJ structure destruction in *Ndrg2^-/-^* mice.

**Figure 4.**
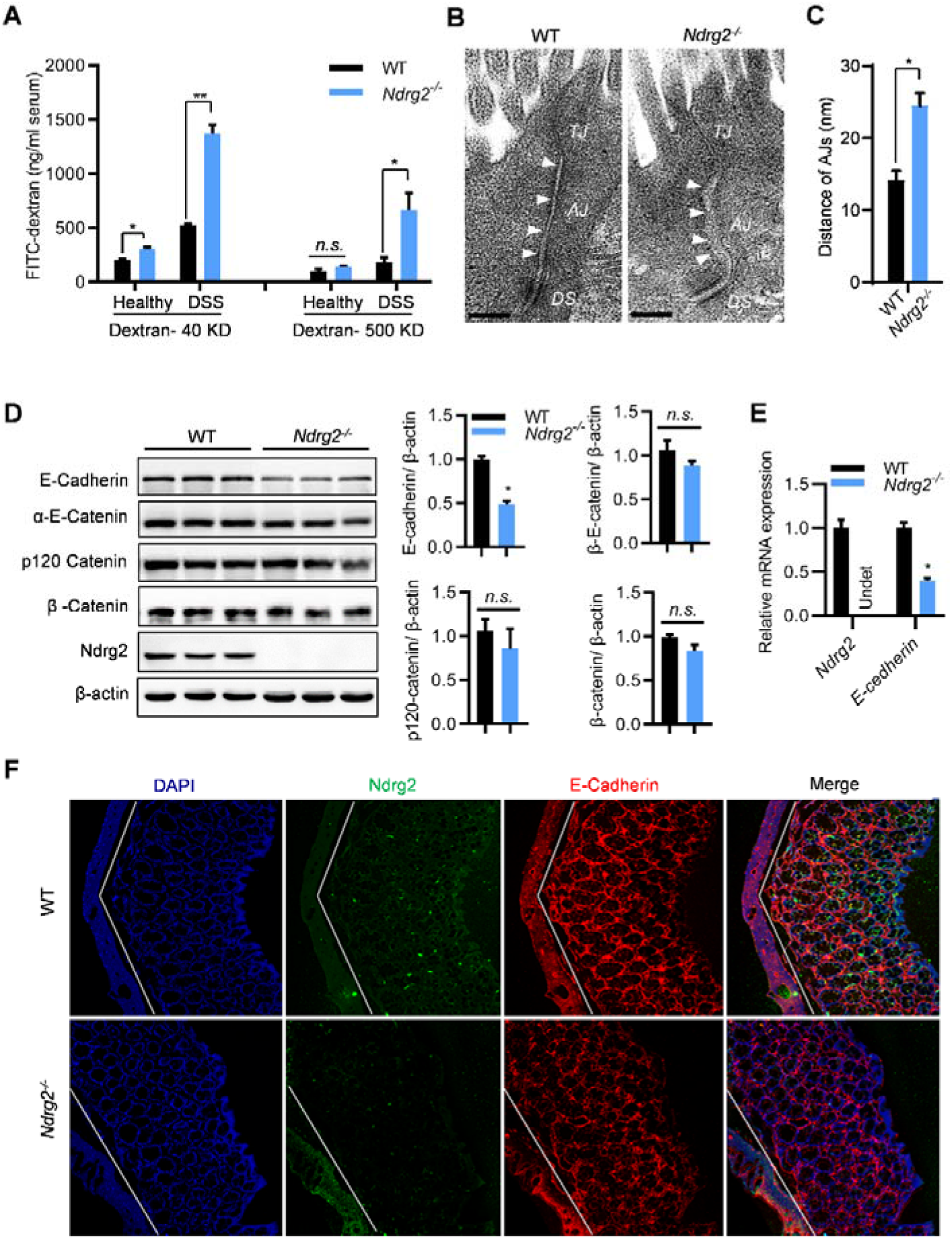
*Ndrg2* deletion disrupted adherens junctions (AJs) integrity in the colonic epithelium. **(A)** Intestinal permeability was measured by determining serum FITC-dextran concentrations (40 KD and 500 KD respectively) (n=8/group). **(B)** AJs structure detection in the colonic epithelium of WT and *Ndrg2^-/-^* mice via electron microscopy. Representative image of 6 mice of each group. Bar: 200 nm. **(C)** Quantification of AJs distances in the colonic epithelium of WT and *Ndrg2^-/-^* mice (n=6/group). **(D)** Western blotting detection for E-Cadherin, α-E-Cadherin, p120 Cadherin, β-Catenin, Snail and Ndrg2 expression in the IECs of WT and *Ndrg2^-/-^* mice (left panel, n=3/group), and band intensity quantification on Right panel. **(E)** qRT-PCR analysis of E-cadherin. **(F)** Immunofluorescence micrographs for NDRG2 (green) and E-cadherin (red) expression in the colons of WT and *Ndrg2^-/-^* mice (n=6/group). **(A, C, D, E)** Data are represented as the mean ± SEM. *p<0.05, **p<0.01, n.s., no significance.

The core components of AJ include E-cadherin and the catenin family members, p120-catenin, β-catenin and α-catenin(Hartsock and Nelson, 2008). We are eager to know the reasons of AJ destruction in the *Ndrg2^-/-^* epithelium. As shown in figure 4D, there was no obvious alteration of p120-catenin, β-catenin and α-catenin expression in *Ndrg2^-/-^* mice (figure 4D). However, intestinal *Ndrg2* loss caused dramatic decrease of E-cadherin both in the protein and mRNA levels (figure 4D, E), suggesting the transcriptional regulation of E-cadherin by NDRG2. Coincidently, immunofluorescence analysis showed that E-cadherin was significantly decreased in the colonic epithelium of *Ndrg2^-/-^* mice (figure 4F), suggesting that *Ndrg2* loss resulted in E-cadherin down-regulation in vivo. Remarkably, E-cadherin attenuation in *Ndrg2^-/-^* mice was much more obviously following spontaneous colitis with ageing (Supplementary figure 5A, B). These findings demonstrated that *Ndrg2* deficiency disrupts AJ structure integrity in the colonic epithelium most likely by suppressing E-cadherin expression, thus increasing colonic permeability and inflammation.

### NDRG2 augments E-cadherin expression via promoting the ubiquitylation and degradation of Snail

The aforementioned results prompted us to uncover how *Ndrg2* loss caused the transcriptional repression of E-cadherin. However, the current studies didn’t support the concept of NDRG2 as a transcriptional regulator per se(Li et al., 2017; Nakahata et al., 2014; Shen et al., 2018). It’s reasonable for us to further analyze whether *Ndrg2* deficiency alters the expression of key transcriptional regulators of E-cadherin, such as Snail, Slug and ZEB1/2 etc(Krebs et al., 2017; Wang et al., 2013). Notably, Snail was one of the most changed regulators and increased significantly in *Ndrg2* deficient IECs and mouse embryonic fibroblast (MEF) cells (figure 5A). We further confirmed the alteration of Snail and E-cadherin in human colorectal cancer cells with NDRG2 knockdown (Supplementary figure 6A, B). Nextly, by subjecting different FLAG-tagged NDRG2 truncations (figure 5B, right panel)(Shen et al., 2018), we found Snail expression was significantly decreased with full length NDRG2 (NDRG2/FL), but obviously abolished by NDRG2 N-terminal deletion (NDRG2/ΔN) (figure 5B, left panel), suggesting the key role of N-terminal of NDRG2 for Snail regulation. With chase assay of protein stability, we noticed NDRG2/FL could obviously shorten the half-life of Snail protein (figure 5C, D). On the contrary, protein stability of Snail was significantly increased in *Ndrg2* deficient MEF cells (figure 5E). However, the Snail suppression by NDRG2 was almost completely reversed by proteasome inhibitor MG132 (figure 5F), strongly supporting NDRG2 might promote Snail protein degradation through ubiquitin-proteosome pathway.

**Figure 5.**
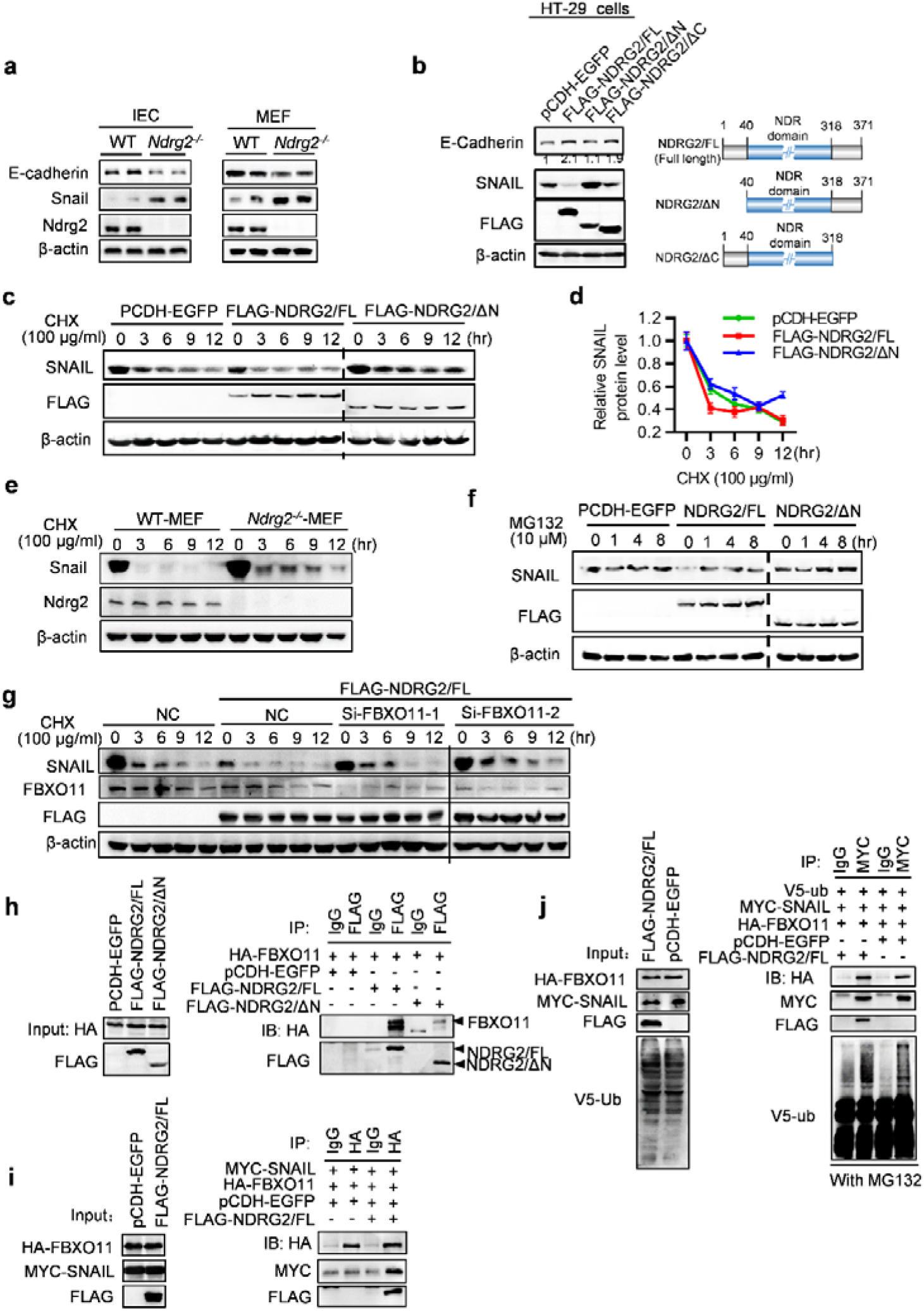
NDRG2 augmented E-cadherin expression *via* promoting the ubiquitylation and degradation of SNAIL. **(A)** Western blotting analysis of E-cadherin, Snail and Ndrg2 protein levels in intestinal epithelial cells (IECs) separated from WT and *Ndrg2^-/-^* mice (Left panel), and also in mouse embryonic fibroblast (MEF) cells established from WT and CMV-Cre; *Ndrg2^-/-^* mice (Right panel). **(B)** Western blotting analysis of endogenous E-Cadherin and SNAIL protein levels in full length NDRG2 (NDRG2/FL), N-terminal truncation (NDRG2/ΔN) and C-terminal truncation (NDRG2/ΔC) stably overexpressing HT-29 cells (Left panel); Graphical representation of NDRG2/FL, NDRG2/ΔN and NDRG2/ΔC truncation mutants (Right panel). **(C)** With 100 μg/ml cycloheximide (CHX) treatment, western blotting detection for SNAIL protein half-life in the cells mentioned above, and quantification in **(D)** by ImageJ software. **(E)** Western blotting detection of Ndrg2 and Snail protein levels in MEF cells as mentioned above treated by CHX for indicated time. **(F)** Endogenous SNAIL protein expression level in NDRG2/FL and NDRG2/ΔN stably overexpressing HT-29 cells treated with 10 μM MG132 for indicated time. **(G)** Using CHX (100 μg/ml) treatment and two siRNAs targeting to FBXO11 with NC as negative control, western blotting analysis of SNAIL, FBXO11 and FLAG-NDRG2 protein expression in parental or FLAG-NDRG2/FL overexpressing SW480 cells. **(H)** HEK293T cells were co-transfected with plasmids expressing HA-FBXO11, FLAG-NDRG2/FL and FLAG-NDRG2/ΔN as indicated respectively, with EGFP as control. The cell lysate was immunoprecipitated with FLAG antibody or IgG with anti-HA and anti-FLAG as primary antibodies for western blotting analysis. **(I)** HEK293T cells were co-transfected with plasmids as indicated respectively. Cell lysates were immunoprecipitated with anti-HA antibody with anti-Myc/FLAG antibodies for western blotting analysis. **(J)** HEK293T cells were co-transfected with plasmids expressing HA-FBXO11, Myc-Snail and V5-Ubiquitin together, with either EGFP or FLAG-NDRG2/FL. Cells were treated with 10 μM MG132 for 6 hrs before cell lysates were immunoprecipitated with anti-FLAG antibody, and detected the poly-ubiquitylated Snail protein with anti-V5 antibody. All the experiment were repeated at least 3 times independently.

There are several reported E3 ligases responsible for Snail ubiquitylation, including β-TRCP1/Fbxw1, Fbxo11, Fbxl14 and Fbxl5 etc(Diaz and de Herreros, 2016; Zheng et al., 2014). So it’s curious for us to know which E3 ligase is responsible for Snail ubiquitylation mediated by NDRG2. Using siRNA strategy to knockdown these E3 ligases respectively and compare the protein stability of Snail in NDRG2 overexpression cells (data not shown). Remarkably, siRNA targeting FBXO11 could obviously rescue the Snail protein stability inhibited by NDRG2 (figure 5G, Supplementary figure 7). Based on this finding, we hypothesized that NDRG2 could enhance the interaction of FBXO11 and Snail to promote the ubiquitylation and degradation of Snail. Immunofluorescence assay demonstrated the cellular co-localization of NDRG2 with Snail and FBOX11 respectively (Supplementary figure 8). Also as the immunoprecipitation assay shown in figure 5H, NDRG2/FL could obviously interact with FBXO11 and significantly strengthen FBXO11 targeting to Snail (figure 5I) to further promote Snail ubiquitination (figure 5J). Our data demonstrated that NDRG2 could enhance E3 ligase FBXO11 with Snail to induce ubiquitin-mediated degradation of Snail, finally augmented E-cadherin expression.

### *Ndrg2* loss in intestinal epithelial cells increases colitis-associated colorectal cancer

To further evaluate the role of *Ndrg2* in colitis-associated tumor development, mice were subjected to Azoxymethane (AOM) intraperitoneal injection followed with three cycle of DSS-H_2_O rotation treatment (figure 6A). Then the mice were sacrificed and colon tissues were collected. After the treatment of AOM/DSS, both WT and *Ndrg2^-/-^* mice developed colon tumors mainly in the distal to middle colon, and the *Ndrg2^-/-^* mice showed obvious increased tumor numbers and size compared with the WT (figure 6B). This phenotype was confirmed by HE staining as typical cancerization was observed on distal colon tissue slides (figure 6C). To further analyze the carcinoma formation in WT and *Ndrg2^-/-^* mice, we calculated the distribution, number and size of tumors and observed significant increasing of tumors located in the middle and terminal colon (figure 6D, E, F). Meanwhile, the number of tumors with all volume scales (under 2 mm, 2-4 mm and above 4 mm) were increased in *Ndrg2^-/-^* mice (figure 6G, H, I). Our data clearly demonstrated that intestinal *Ndrg2* loss increased susceptibility to colitis-associated tumor development.

**Figure 6.**
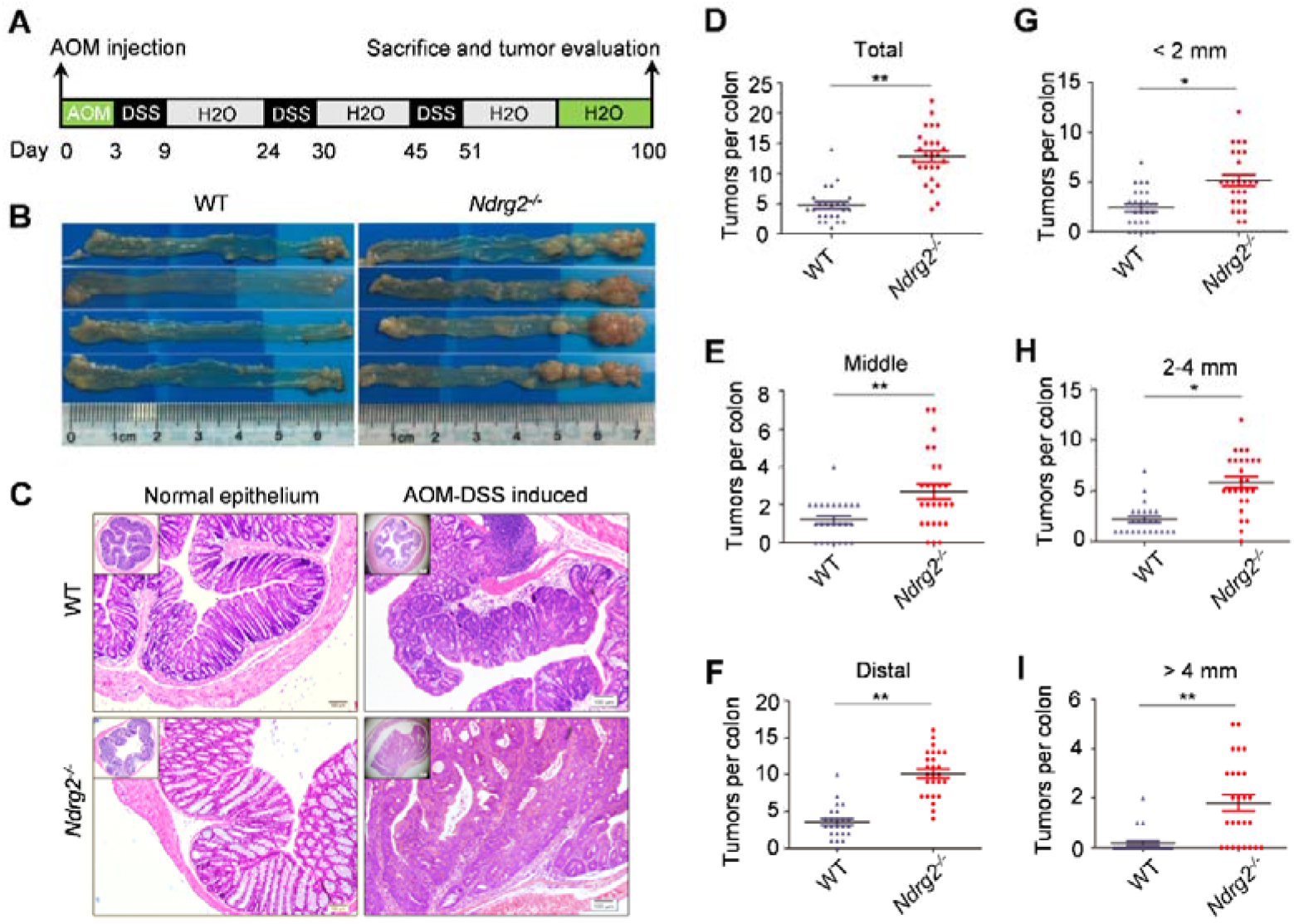
Intestinal deletion of *Ndrg2* enhanced the initiation and progression of AOM-DSS induced colorectal carcinoma. **(A)** Method of the treatment of AOM-DSS in our study. **(B)** Tumor bearing colon sections from WT and *Ndrg2^-/-^* mice after AOM-DSS treatment. **(C)** HE staining of normal epithelium and AOM-DSS induced colorectal carcinoma in WT and *Ndrg2^-/-^* mice. Bar: 100 μm. **(D-F)** Quantification analysis of tumor numbers from the whole, middle and distal colons between WT and *Ndrg2^-/-^* mice respectively. **(G-I)** Quantification analysis of tumor numbers with small size (<2 mm), middle size (2-4 mm) and big size (>4 mm) from the whole colon between WT and *Ndrg2^-/-^* mice respectively. WT: n=24; *Ndrg2^-/-^*: n=26. *p<0.05, **p<0.01.

### NDRG2 down-regulation attenuates E-cadherin expression and enhances inflammation in human IBD patients

To confirm the role of NDRG2 in clinic IBDs initiation, we further examined whether NDRG2 affected inflammation in human UC and CD patients (Supplementary Table 3). Interestingly, in UC patients, NDRG2 expression was significantly positively correlated with E-cadherin, and negatively correlated with Snail (figure 7A, B, C). Most importantly, NDRG2 expression was negatively correlated with inflammation levels and CD68^+^ macrophage recruitment. Higher NDRG2 expression was correlated with reduced inflammation and CD68^+^ macrophage recruitment at inflammatory foci, while decreases in NDRG2 expression enhanced inflammation and CD68^+^ macrophage infiltration (figure 7A, D, E).

**Figure 7.**
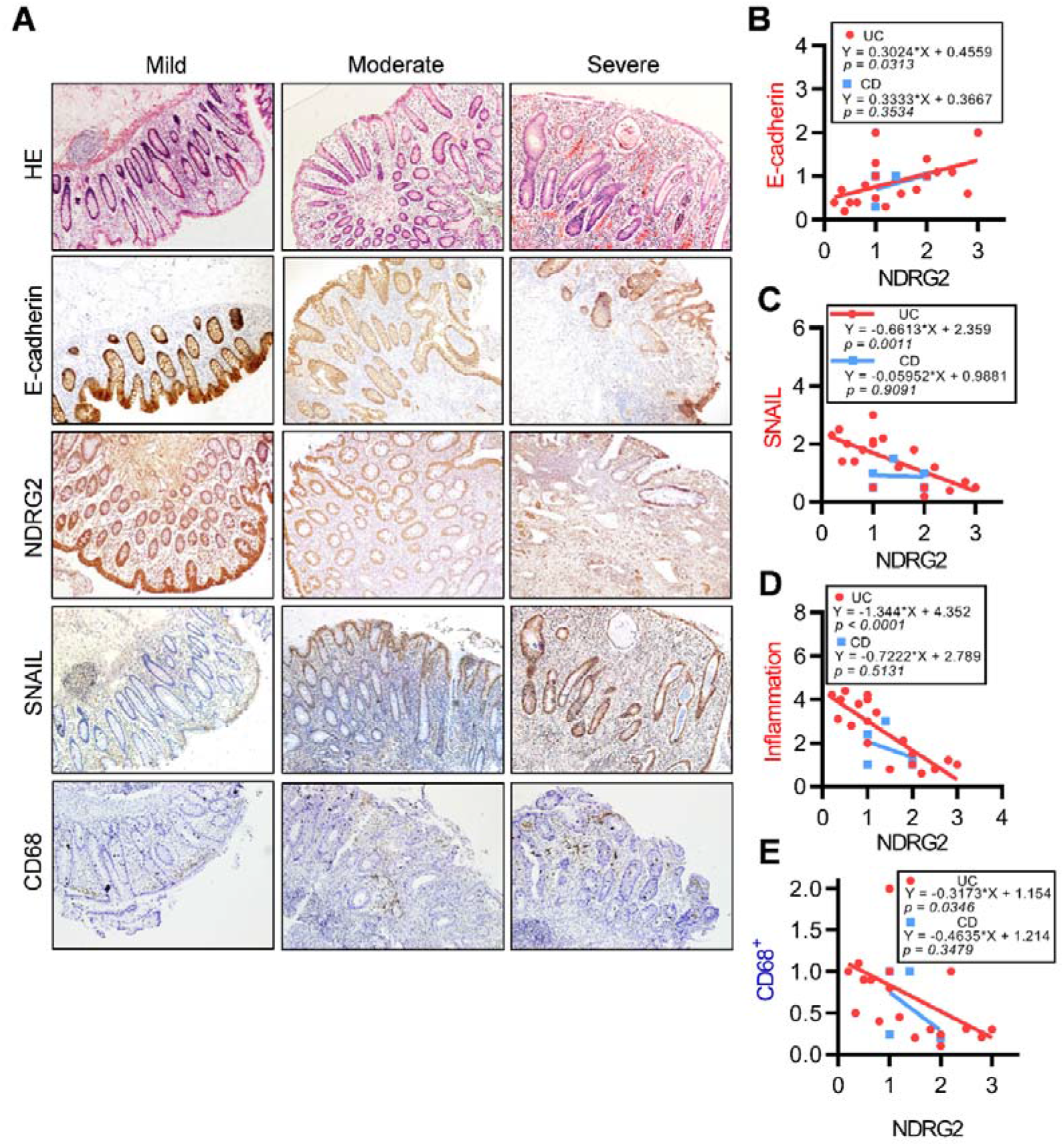
NDRG2 positively correlated with E-cadherin and negatively correlated with Snail and inflammation in human IBD patients. **(A)** Immunohistochemistry of NDRG2, E-cadherin, Snail, CD68^+^ and HE staining on human UC tissues with mild, moderate and sever inflammation degree. Bar: 100 μm. **(B-E)** Linear regression and Pearson’s correlation analysis of NDRG2 with inflammation degree, E-cadherin and Snail in UC and CD samples respectively.

Taken together, our findings demonstrated that NDRG2 could positively enhance E-cadherin expression to maintain AJ structure and integrity of intestinal epithelial barrier, thus suppressing colonic inflammation. However, intestinal *Ndrg2* loss led to E-cadherin attenuation and AJ structure destruction, resulting in increase of intestinal epithelial permeability, thus promoting colitis and colitis-associated tumor development (figure 8).

**Figure 8.**
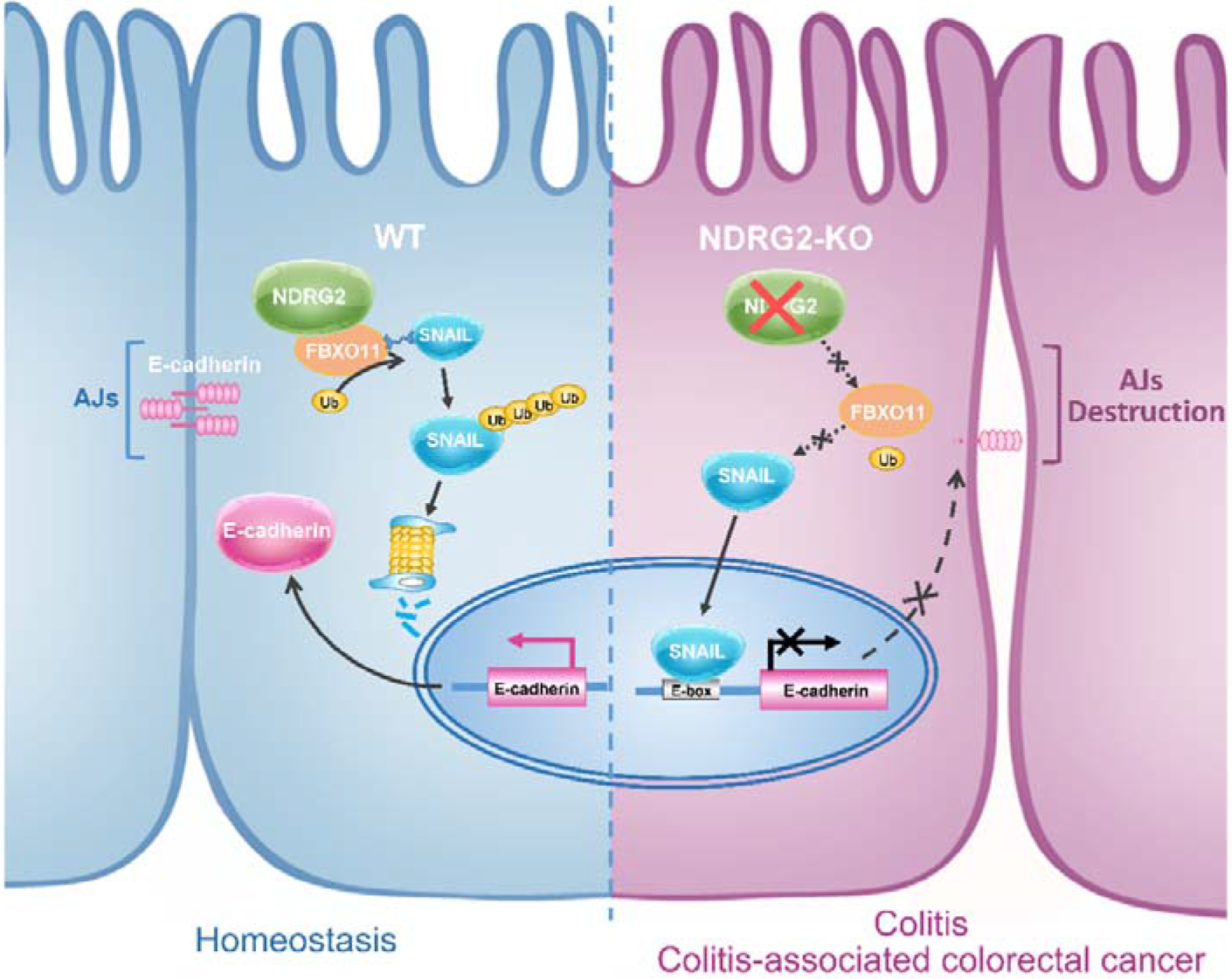
A schematic for the role of NDRG2 regulating Snail, E-cadherin and adherens junction structure integrity in colitis and colitis-associated tumor development. Intestinal *Ndrg2* loss disrupts AJs structure in the colonic epithelium by suppressing E-cadherin expression via abrogating Snail ubiquitination and degradation through E3 ligase FBXO11-dependent signaling, thus increasing colonic permeability, colitis and colitis-associated tumor development in *Ndrg2^-/-^* mice.

## DISCUSSION

The gastrointestinal tract (GI) is chronically exposed to large numbers of foreign antigens, toxic molecules and microorganisms, and the epithelial barrier makes important contributions to GI health(Peterson and Artis, 2014; Taddei et al., 2008). Epithelial barrier dysfunction is believed to contribute to IBD initiation and pathogenesis, mainly by damaging the structure of tight junctions (TJs) and/or adherens junctions (AJs). Previous reports have demonstrated that AJ and TJ disruption is associated with progressive colonic inflammation in human UC and CD due to increased permeability of LPS, peptidoglycan and N-formyl-L-methionyl-L-leucyl-L-phenylalanine (fMLP), which diffuse from the gut lumen to the lamina propria(Kobayashi et al., 2013; Landy et al., 2016).

Our study explored for the role of NDRG2 in intestinal inflammation pathogenesis and determined that NDRG2 modulated AJ structure *via* FBXO11/Snail/E-cadherin signaling regulation *in vivo*. NDRG2 was initially discovered and demonstrated to be a tumor suppressor gene by our group. We found that NDRG2 played roles in cancer cell proliferation, differentiation and invasion(Li et al., 2018; Yao et al., 2008). NDRG2 expression was positively correlated with cancer prognosis and overall survival(Chu et al., 2011). Furthermore, NDRG2 regulated EMT by controlling E-cadherin expression and participated in TGF-β-induced oncogenesis in late-stage CRC(Kim et al., 2013; Shen et al., 2014).

E-cadherin is an important cytomembrane component and controls cell-cell adhesion by conjugating with β-catenin, α-catenin and p120-catenin complexes to modulate the spaces between epithelial cells(Bulgakova and Brown, 2016; Orsulic et al., 1999). Previous studies confirmed that reduced E-cadherin aggravated DSS-induced colitis(Grill et al., 2015; Wheeler et al., 2001). Our work demonstrated that *Ndrg2* loss in intestinal epithelial cells caused AJ destruction but did not disrupt tight junction structure, as determined *via* electron microscopic analysis. Moreover, we observed the obvious reduced E-cadherin expression with no significant alterations of β-catenin or p120-catenin in *Ndrg2^-/-^* mice, indicating that AJ destruction caused by *Ndrg2* loss is mainly induced by E-cadherin suppression. Therefore, intestinal *Ndrg2* deletion decreased E-cadherin expression and destroyed the integrity of adherens junctions to increase the epithelial barrier permeability, resulting in spontaneous colitis with ageing, also increasing the susceptibility to DSS and TNBS-induced colitis. Noticeably, we further found that intestinal *Ndrg2* loss could obviously promote the colitis-associated tumor development, which is highly coincident with its tumor suppressor function.

Snail and Slug are believed to be the most important transcriptional repressors of E-cadherin in various types of solid tumors and tissues(Hajra et al., 2002; Yokoyama et al., 2001). Our data showed that Snail significantly increased in *Ndrg2* deficient IECs and MEF cells, suggesting reduced E-cadherin expression by *Ndrg2* loss might be mainly through Snail upregulation. Furthermore, we firstly demonstrated that NDRG2 could strengthen the interaction of E3 ligase FBXO11 with Snail to promote the ubiquitination and degradation of Snail. Thus, as shown in figure 4 and figure 5, intestinal *Ndrg2* deletion disrupts adherens junction structure in the intestinal epithelium by suppressing E-cadherin expression via promoting FBXO11-Snail interaction for Snail ubiquitination and degradation, thus increasing colonic permeability, colitis and colitis-associated tumor development.

Inflammatory cell infiltration plays an important role in IBD(Koelink et al., 2014), and we found that *Ndrg2* loss facilitated inflammatory cell recruitment and reduced NDRG2 expression was associated with the severe inflammatory state in clinical samples of UC patients. Most importantly, we confirmed the positive correlation of NDRG2 with E-cadherin, and negative correlation with Snail in UC samples. However, it’s hardly to recognize this similar pattern in CD patients, partially due to the limited CD patient samples, which might also suggest the different function of NDRG2 in UC and CD. *Ndrg2* loss-induced E-cadherin repression leads to increase of intestinal permeability and pro-inflammatory cytokine production, thereby triggering more immune response to further aggravate colitis progression.

In summary, our work confirmed that NDRG2 played important role in colitis initiation and pathogenesis by enhancing E-cadherin expression and regulating AJ structure integrity to keep gut homeostasis. However, intestinal *Ndrg2* deficiency resulted in attenuated E-cadherin expression and AJ structure disruption, finally promoting the colitis and colitis-associated tumor development (figure 8). Thus, NDRG2 might be an important diagnosis and prognosis marker of colitis and colitis-associated colorectal cancer.

## MATERIALS AND METHODS

Experimental methods, including statistical analysis are described in detail in the online supplementary information.

## Acknowledgments

This work was supported by the National Natural Science Foundation of China (No. 81770523, 31571437, 81672751), the Founds for Creative Research Groups of China (No. 81421003), and the State Key Laboratory of Cancer Biology Project (CBSKL201406, CBSKL2017Z08 and CBSKL2017Z11).

## Author contributions

Design of studies: JZ, JPL, LBY and KCW. Performance of experiments: MYW, YZM, LLS, YQX, ZHG, LJL, XB, HYQ Clinical assessments: YQX, ZSL, ZW,. Writing of manuscript: JZ, MYW, YZM. All authors approved the manuscript.

