## Supplementary Figure legends for "NDRG2 Regulates Adherens Junction Integrity to Restrict Colitis and Tumorigenesis" (from another version of this preprint: version 2, 2019-11-27)

**Supplementary Figure 1. Characteristics of intestine-specific conditional *Ndrg2* knockout mice (*Ndrg2*^-/-^). (A)** Construction of the targeting vector and the wild-type allele, floxed allele and deleted allele of the mouse *Ndrg2* gene. *Loxp* fragments were placed on both sides of exons 2 - 6 to delete them, and the *neo^r^* cassette was identified for vector selection. **(B)** Western blot detection of Ndrg2 protein expression in brain, heart and liver tissue samples from WT and *Ndrg2*^-/-^ mice. **(C)** Western blot detection of Ndrg2 protein expression in total colon tissue and IECs from WT and *Ndrg2*^-/-^ mice respectively (n=2/group).

**Supplementary Figure 2. *Ndrg2* deficiency enhanced TNBS-induced colitis.**

**(A)** WT and *Ndrg2*^-/-^ mice was slowly instilled with 100 μl of 3 mg TNBS (dissolved in 50% ethanol) into colon to induce colitis (n=8/group). Body weight was monitored from Day 1 to Day 6. Data are represented as the mean ± SEM. *p<0.05. **(B)** The colon morphology of TNBS-induced colitis in WT and *Ndrg2*^-/-^ mice.

**Supplementary Figure 3. *Ndrg2* deficiency enhanced inflammation in mice colon section. (A)** WT and *Ndrg2*^-/-^ mice were treated with 2.5% DSS (n=12/group). Flow cytometry analysis was performed on days 0 and 10 after DSS treatment. CD11b^+^ stained monocytes, CD11b^+^ F4/80^+^ stained macrophages, CD11b^+^ Gr-1^+^ stained neutrophils, and CD11b^+^ Siglec-F^+^ stained eosinophils. **(B)** qRT-PCR for the mRNA expression of indicated chemokines and cytokines. Data are represented as the mean ± SEM. *p<0.05, **p<0.01, n.s., no significance (Student’s t test).

**Supplementary Figure 4.** **LPS induced inflammation in the colonic epithelium in a dose-dependent manner.** WT mice were treated with ampicillin for 1 week before LPS administration, and then LPS was injected into the gut lumen at doses of 1 mg/kg and 0.1 mg/kg per mouse respectively. Intestinal epithelial cells (IECs) were isolated, and total RNA was extracted according to the manufacturer’s instructions. **(A-E)** qRT-PCR for the mRNA expression of *IL-1β, IL-6, TNF-α, CXCL-1,* and *CXCL-2* expression (n=4). Data are represented as the mean ± SEM. *p<0.05.

**Supplementary Figure 5. Intestinal *Ndrg*2 loss caused obvious E-cadherin attenuation in ageing mice with spontaneous colitis.** **(A)** (Top) Schematic diagram for analyzing the occurrence of spontaneous colitis in 8 weeks age WT and *Ndrg2^-/-^* mice after withdrawal of ampicillin treatment for 1 week. (Bottom) Western blotting analysis of endogenous E-cadherin protein level in IECs from mice of WT and *Ndrg2^-/-^* mice with specific-pathogen-free (SPF) feeding condition. **(B)** (Top) Schematic diagram for analyzing the occurrence of spontaneous colitis in 36 weeks age WT and *Ndrg2^-/-^* mice after withdrawal of ampicillin treatment for 1 week. (Bottom) Western blotting analysis of endogenous E-cadherin protein level in the intestinal epithelial cells (IECs) of WT and *Ndrg2^-/-^* mice with specific-pathogen-free (SPF) feeding condition. Representative results of 2 mice in each group.

**Supplementary Figure 6.** **NDRG2 knockdown caused the alteration of E-cadherin and Snail in HT-29 and Caco-2 cells.** Western blotting for E-cadherin, NDRG2 and Snail expression in HT-29 cells **(A)** and Caco-2 cells **(B)** with NDRG2 siRNA knockdown.

**Supplementary Figure 7.** SW480 cells were transfected using two siRNAs targeting to FBXO11 with NC as control. Western blotting was employed for the detection of endogenous SNAIL, FBXO11 and NDRG2 protein expression levels with MG132 (10 µM) treated for indicated time.

**Supplementary Figure 8. NDRG2 co-localized with FBXO11 and SNAIL.** FLAG-NDRG2/FL or FLAG-NDRG2/ΔN were co-transfected with HA-FBXO11 or MYC-SNAIL into HEK293T cells respectively, empty vector were served as control. After transfection for 24 hours, cells were fixed and then subjected to Immunofluorescence with anti-HA antibody allowed FBXO11 visualization, anti-FLAG antibody represented NDRG2/FL or NDRG2/ΔN, and anti-MYC antibody indicated SNAIL. Hoechst 33258 was used to visualize nuclei. Scale bars indicate 10 μm.
