## Supplementary Methods for "NDRG2 Regulates Adherens Junction Integrity to Restrict Colitis and Tumorigenesis" (from another version of this preprint: version 2, 2019-11-27)

**Mice**

Intestine-specific *Ndrg2* knockout mice (*Ndrg2*^-/-^) were generated by Shanghai Biomodel Organism Science & Technology Development Co., Ltd and maintained on a C57BL6/J background. All animals were raised under specific-pathogen-free conditions. The wild-type (WT) and *Ndrg2*^-/-^ mice were age-matched, and all the mice that were used in the experiments were male. Typically, at least 8 mice were included each treatment group, and all experiments were repeated at least three times.

**Spontaneous murine colitis analysis**

To analyze spontaneous colitis development, 8 weeks and 36 weeks age *Ndrg2*^-/-^ mice were first given ampicillin (1 g/L) water for one week, and then replaced with normal water. The animals were raised under specific-pathogen-free conditions and were sacrificed at indicated time, and colon tissues were collected for further detection. Age-matched WT mice were used as control. Throughout the experiment mice were free access to food and water.

**DSS-induced colitis and colitis-associated colorectal cancer**

For DSS-induce colitis model, 8 weeks age male mice were treated with 2.5% DSS (w/v, MW 36 - 50 kDa; MP Biomedicals, Solon, OH) in drinking water for 5 days, followed by DSS-free water for another 5 days. Before sacrificing the mice and evaluating their inflammatory responses. Distal colon were obtained for histological analysis, and IECs and LPCs were separated with the total colon.

To induce the colitis-associated colorectal cancer, we used the combination of carcinogen Azoxymethane (AOM) with repeated treatment of DSS in drinking water. Mice were treated with intraperitoneally injection with a single dose of AOM (12 mg/kg). After 3 days, 2% DSS dissolving in the drinking water were given for 6 days (the DSS solution was refreshed at day 3), followed by 15 days of regular drinking water. The DSS treatment was repeated for another two cycles. Mice were sacrificed 100 days after AOM injection, colons were removed and the colon tumors were dissected for further analysis.

**TNBS-induced murine colitis**

The 2, 4, 6-Trinitrobenzenesulfonic acid (TNBS)-induced experimental colitis was performed in C57BL/6 mice (B6) as previously described. Briefly, *Ndrg2*^-/-^ mice and their littermate controls were anesthetized after fasting for 12 hours, then 100 μl of 3 mg TNBS (dissolved in 50% ethanol) was slowly instilled into colon using a 3.5 F catheter which was inserted intra-rectally 4 cm from the anus. Mice were held in an inverted vertical position for 30 s after instillation to ensure even distribution of TNBS throughout the entire colon and cecum. Mice from respective control groups received PBS in a comparable volume by the same route.

**Histological score determination**

Distal colonic tissue segments were fixed in 4% neutral-buffered formalin, embedded in paraffin, cut into 8 μm-thick sections, and stained with H&E to histopathologically detect cell hyperplasia, epithelial damage, and inflammatory cell infiltration and to perform histological scoring, which was completed using the following scale: 0: no evidence of inflammation; 1: low-level inflammation; 2: moderate inflammation; 3: high-level inflammation and 4: maximal inflammation.

**Isolation of intestinal epithelial cells (IECs) and lamina propria cells (LPCs)**

Mice were sacrificed and total colon tissues were dissected, opened longitudinally and washed extensively in cold PBS. Cut the colon tissue into 1 cm segments, and wash the segments with fresh PBS. Incubated the tissues with digestion buffer contained 8 mM EDTA and 1 mM DTT at 37℃ for 20 min, then replaced digestion buffer with cold PBS, shacked vigorously for 1 min. The process was repeated once. Combine the supernatants and centrifuged at 2000 rpm for 5 min at 4℃. Digested the pellet cells with Dispase Ⅱ (0.2 mg/ml, Roche #04942078001) in PBS at 37 min for 5 min. Filtered the suspension with 70 μm and 40 μm nylon filters (BD biosciences), and centrifuged at 2000 rpm for 5 min at 4℃, resuspended the cells in cold PBS. To obtained LPCs, tissues after EDTA incubation were cut into small pieces and digested with the mix of collagenase A (2 mg/ml, Roche #10103586001) and DNase I (0.05 mg/ml, Sigma #D4263) in PBS at 37℃ for 35 min. After incubation, replaced the enzyme mixture with cold PBS, shacked for 2 min. Repeated the process once. The suspension was combined and passaged through 70 μm and 40 μm nylon filters. Centrifuged at 4000 rpm for 5 min at 4℃, and resuspended the pellet cells in cold PBS. IECs were used for extraction of RNA and Western blot analysis. For quantification of immune cells, both IECS and LPCs were combined and stained with CD11b, Gr-1, Siglec-F, F4/80.

**Analysis of intestinal permeability**

Fluorescein isothiocyanate-conjugated (FITC) dextran with average mol wt 40,000 (Sigma-Aldrich #FD40) or with average mol wt 500,000 (Sigma-Aldrich #FD500S) were dissolved at a concentration of 100 mg/ml in PBS respectively. According to the manufacturer’s instructions, all experimental mice was treated with water starvation for at least 10 hours, then FITC-dextran was administered to each mouse (44 mg/100 g body weight) by oral gavage with a needle attached to a 1 ml syringe. After 4 hours, anesthetized the mice with amobarbital and collected at least 400 μl of blood from each mouse. Serum was separated and collected according to the manufacturer’s instructions and stored in the dark at 4℃. Diluted 150 μl of serum with an equal volume of PBS, and added 100 μl of the dilution to 96-well microplate in duplicate. The concentration of FITC-dextran were determined by spectrophoto fluorometry with an excitation of 485 nm (20 nm band width) and an emission wavelength of 528 nm (20 nm band width). Serum from untreated mice was to determine the background.

**Flow cytometry staining**

For detection of monocytes, macrophages and neutrophils, the following fluorochrome-conjugated antibodies were used: Anti-Mouse CD11b FITC (eBioscience #11-0112); Anti-Mouse Ly-6G (Gr-1) PE (eBioscience #12-5931); Anti-Mouse F4/80 Antigen APC (eBioscience #17-4801); PE Rat Anti-Mouse Siglec-F (BD Pharmingen™ #562068). According to the methods reported, CD11b staining was used to detect monocytes, combination of CD11b and Ly-6G (Gr-1) was used to detect neutropihls, and combination of CD11b and Siglec-F was used to quantify eosinophils, while macrophages was detected with CD11b and F4/80. Both IECs and LPCs were combined for quantification of the immune cells. Total cells were resuspended with PBS containing 5% FBS, and at least 10^6^ cells were used for the staining.

**Analysis of adherens junction by transmission electron microscopy**

Mice were perfused through left atrium with mix of 4% paraformaldehyde and 1% glutaraldehyde. Pieces of colon tissue were fixed in 3% glutaraldehyde with 2% OsO4 for 6 hrs at 4℃. Dehydrated the tissue at 4℃ through a series of acetone concentration (50, 70, 90, 96, 100%) and embedded in Epon 812 epoxy resin. Sections were cut with a thickness of 70 nm and collected on 200-mesh, Formvar-coated copper grids. The grids were stained with uranylacetate and lead citrate, micrographs were collected with a JEOL JEM 1230 electron microscope (JEOL, Japan).

**RNA isolation and quantitative real-time PCR (qPCR)**

Total IECs RNA was isolated from healthy and DSS-treated WT and *Ndrg2*^-/-^ mice using Trizol reagent (Takara, Dalian, China), according to the manufacturer’s instructions. Primers sequences for quantitative real-time polymerase chain reaction (qRT-PCR) are listed in Supplementary Table 1 and Table 2. qRT-PCR data were analyzed by the 2^-ΔΔCT^ method, with *GAPDH* and *β*-actin as the housekeeping gene.

**Co-immunoprecipitation assay**

FLAG tagged NDRG2 (full length and truncated form named as NDRG2/FL, NDRG2/ΔN, NDRG2/ΔC) plasmids were co-transfected with Myc-Snail, HA-FBXO11 (gift from Dr. Yibin Kang, Princeton University) and V5-Ubiquitin or indicated plasmids as described in figures in 60 mm dishes using LipoPlus transfection reagent (Sage Creation, Beijing, China) according to the manufacturer's instructions. For ubiquitination assay, cells were treated with 10 μM MG132 for indicated times after 2 days transfection. Then cells were harvested with lysis buffer (Beyotime, P0013), proteins were quantified using Pierce BCA Protein Assay Kit (Thermo Scientific, 23225) and adjusted to equal concentration. Cell lysates were then immunoprecipitated with primary anti-FLAG (Sigma, F1804) or anti-Myc (CST, 66004) or anti-HA (CST, 3724) or anti-V5 (Sigma, V8137) at 4℃ over night. Then 70 μL protein A/G sepharose (Abcam, ab193262) were added and incubated for 3 hrs. After 3 times washing using NETN wash buffer, Tris 20 mM (pH 8.0), NaCl 100 mM, EDTA 1 mM, NP-40 0.5%, sepharose beads were collected and re-suspended with 70 μL 2xSDS buffer. Pull-down samples and with 5% input were resolved on SDS-PAGE gel and transferred to NC membranes. Then the membranes were detected with appropriate antibodies and visualized using Tanon 6200 Luminescent Imaging Workstation.

**Western blotting analysis**

Total proteins were extracted from intestinal epithelial cells (IECs) according to the manufacturer’s recommendations. Sodium dodecyl sulfate polyacrylamide gel electrophoresis (SDS-PAGE) and western blot analysis were performed with 40 μg protein. The following antibodies were used: β-actin (Cell Signaling, #12262), NDRG2 (Cell Signaling, #5667), E-cadherin (Cell Signaling, #3195), Snail (Cell Signaling, #3879), Slug (Cell Signaling, #9585), Claudin-1 (Cell Signaling, #13255).

**Immunofluorescence staining**

For murine studies, 8 μm frozen segments of colon from WT and ΔIEC mice were fixed in cold acetone for 15 min at -20℃, and then 3% BSA was used for blocking at 25℃ for 1 hr. Anti-E-cadherin (1:100) and NDRG2 (1:80) was use for incubation at 4℃ overnight, and Alexa-Fluor 488- or 647- conjugated antibodies (Invitrogen, 1:2000) was used as the secondary antibodies and incubated for 1 hr at 25℃. Incubation of 1 μg/ml of DAPI was followed for 15 min. Fluorescence analysis was performed with a laser scanning confocal microscope A1 (Nikon). For cellular protein distribution detection, cell were seeded in Rat tail collagen coated dishes and transfected with indicated plasmid for 24 hours, and then gently washed with PBS for 5 min and fixed in 4% paraformaldehyde for 30 min. Immunofluorescence Staining was performed as method above. Anti-HA (1:500), anti-FLAG (1:500) and anti-SNAIL (1:200) were used for incubation at 4℃ overnight.

**Tissue samples and immunohistochemistry staining**

This study was approved by the ethics committee of the Fourth Military Medical University. Totally 19 ulcerative colitis cases and 5 Crohn’s disease patient samples were obtained from Xijing Hospital from April 2013 to October 2016. The clinical information was shown in Supplementary Table 3.

**Study approval**

All animal procedures were reviewed and approved by the Animal Care and Use Committee of Fourth Military Medical University. All animals were housed, cared for, and used in compliance with the *Guide for the Care and Use of Laboratory Animals*. All patients in the disease sample cohort provided full consent for the study.

**Statistical analysis**

Differences in parametric data were evaluated using Student’s t test or one-way analysis of variance (ANOVA). *p<0.05 and **p<0.01 were considered statistically significant.
