## Supplementary Tables for "NDRG2 Regulates Adherens Junction Integrity to Restrict Colitis and Tumorigenesis" (from another version of this preprint: version 2, 2019-11-27)

**Supplementary Table 1.** Primers for qRT-PCR analysis (mouse)

| **Gene (mouse)** | **Forward (5’-3’)** | **Reverse (5’-3’)** |
| --- | --- | --- |
| **GAPDH** | CTTCACCACCATGGAGGAGGC | GGCATGGACTGTGGTCATGAG |
| **IL-6** | AGTTGCCTTCTTGGGACTGA | CAGAATTGCCATTGCACAAC |
| **IL-1β** | GAGAGCCGGGTGACAGTATC | TGACAAACTTCTGCCTGACG |
| **TNF-α** | CGTCAGCCGATTTGCTATCT | CGGACTCCGCAAAGTCTAAG |
| **IL-10** | CCAAGCCTTATCGGAAATGA | TTTTCACAGGGGAGAAATCG |
| **IL-11** | TGTTCTCCTAACCCGATCCCT | CAGGAAGCTGCAAAGATCCCA |
| **IL-17** | GCCCTCAGACTACCTCAACC | ACACCCACCAGCATCTTCTC |
| **IL-12** | AGGTCACACTGGACCAAAGG | TGGTTTGATGATGTCCCTGA |
| **IL-22** | GCAATCAGCTCAGCTCCTGT | CGCCTTGATCTCTCCACTCT |
| **TGFβ1** | GGAGGTACCGCCCGGCCCGC | GACAGCAATGGGGGTTCGGG |
| **CXCL-1** | ACTGCACCCAAACCGAAGTC | TGGGGACACCTTTTAGCATCTT |
| **CCL-5** | GCTGCTTTGCCTACCTCTCC | TCGAGTGACAAACACGACTGC |
| **CXCL-2** | CTGCGCCCAAACCGAAGTCATA | TTCAGGAACAGCCACCAATAAGC |
| **ZO-1** | GCCGCTAAGAGCACAGCAA | GCCCTCCTTTTAACACATCAGA |
| **Claudin-1** | TGCCCCAGTGGAAGATTTACT | CTTTGCGAAACGCAGGACAT |
| **Snail** | CACACGCTGCCTTGTGTCT | GGTCAGCAAAAGCACGGTT |
| **Slug** | CAGCGAACTGGACACACACA | ATAGGGCTGTATGCTCCCGAG |
| **E-cadherin** | CAGTTCCGAGGTCTACACCTT | TGAATCGGGAGTCTTCCGAAAA |

**Supplementary Table 2.** Primers for qRT-PCR Analysis (human)

| **Gene (human)** | **Forward (5’-3’)** | **Reverse (5’-3’)** |
| --- | --- | --- |
| **β-actin** | CATGTACGTTGCTATCCAGGC | CTCCTTAATGTCACGCACGAT |
| **Claudin-1** | CCTCCTGGGAGTGATAGCAAT | GGCAACTAAAATAGCCAGACCT |
| **Snail** | TCGGAAGCCTAACTACAGCGA | AGATGAGCATTGGCAGCGAG |
| **Slug** | TGTGACAAGGAATATGTGAGCC | TGAGCCCTCAGATTTGACCTG |
| **E-cadherin** | CGAGAGCTACACGTTCACGG | GGGTGTCGAGGGAAAAATAGG |

**Supplementary Table 3**. Clinical Information of Ulcerative Colitis Patients

| **Age** | **Gender** | **type** | **position** | **Degree of inflammation** |
| --- | --- | --- | --- | --- |
| 50 | Male | UC | Whole colon | Severe |
| 49 | Female | UC | Transverse colon | Severe |
| 42 | Male | UC | Whole colon | Severe |
| 41 | Male | UC | Whole colon | Severe |
| 24 | Male | UC | Whole colon, rectum | Severe |
| 38 | Female | UC | whole colon | Severe |
| 44 | Male | UC | Part of enteric canal | Severe |
| 41 | Male | UC | Rectosigmoid | Severe |
| 49 | Female | UC | Descending colon, rectum | Severe |
| 47 | Female | UC | Whole colon | Moderate |
| 22 | Male | UC | Whole colon | Moderate |
| 48 | Male | UC | Whole colon | Moderate |
| 22 | Male | UC | Whole colon, distal ileum | Mild |
| 48 | Female | UC | Whole colon, rectum | Mild |
| 41 | Male | UC | Rectum | Mild |
| 27 | Male | UC | Sigmoid | Mild |
| 69 | Female | UC | Distal stomach | Mild |
| 47 | Female | UC | Rectosigmoid | Mild |
| 21 | Male | UC | Whole colon | Mild |
| 19 | Male | CD | Ileocecus | Moderate |
| 60 | Male | CD | Ileocecus | Moderate |
| 26 | Male | CD | Ileocecus | Mild |
| 47 | Male | CD | 75 cm from the anus | Mild |
| 45 | Male | CD | Ileocecus | Mild |
